# Genomic Insights into a Versatile Deep-Sea Methanotroph Constituting the Rare Biosphere of a Brazilian Carbonate Mound Complex

**DOI:** 10.1101/2025.06.09.658676

**Authors:** Ana Carolina de Araújo Butarelli, Fernanda Mancini Nakamura, Francielli Vilela Peres, Flúvio Modolon da Silva, Amanda Gonçalves Bendia, Raissa Basti, Michel Michaelovitch de Mahiques, Paulo Yukio Gomes Sumida, Vivian Helena Pellizari

## Abstract

Recent discoveries of aerobic methanotrophs in non-seep carbonate-rich environments in the deep sea suggest that these organisms may persist as part of the rare biosphere. Recovering rare, active methanotrophs through targeted culturing is essential for understanding their persistence under the oligotrophic non-seep conditions, and for uncovering their genomic adaptations related to the survival in energy-limited ecosystems. In our study, using metagenomic analysis of enrichment cultures from the Alpha Crucis Carbonate Ridge, we discovered *Methylotuvimicrobium crucis* sp. nov., a novel methanotroph representing the rare biosphere in native sediments. Phylogenomic analysis revealed <95% ANI to described species, with genomic evidence of deep-sea specialization including: (1) stress adaptation through cold-shock proteins (*CspA*) and DNA repair systems (*UvrD/LexA*), (2) metabolic versatility via complete methane oxidation *(pmoABC)*, nitrogen fixation (*nifHDK*), and sulfur cycling (*sox/sqr*) pathways, and (3) niche partitioning through biofilm formation (*GGDEF/EAL*) and heavy metal resistance (*CopZ/CzcD)*. Comparative genomics identified a 1,234-gene deep-sea core shared with *M*. sp. wino1, enriched in mobile elements (*TnpA*, prophages) suggesting horizontal gene transfer drives adaptation. While undetected *in situ* amplicon surveys, *M. crucis* exhibited rapid enrichment under methane availability, demonstrating its role as a latent methane filter. These findings contribute for the understanding of the ecological significance of aerobic methanotrophs in deep-sea systems, revealing how rare microbial taxa with genomic plasticity have the potential to influence biogeochemical cycling in deep carbonate-rich environments.

## Introduction

Deep-sea geomorphic structures, including carbonate mounds, pockmarks, and mud volcanoes, serve as archives of past or intermittent fluid and gas seepage, shaping distinct ecosystems in regions such as the Southwest Atlantic (1). These environments are characterized by dynamic physicochemical conditions, marked by fluctuating hydrocarbon fluxes and heterogeneous substrates, which drive microbial niche differentiation (2); (3). Notably, carbonate mounds (authigenic formations often linked to microbial activity) act as long-term carbon sinks while harboring microbial consortia critical to biogeochemical cycling (4). Despite their ecological significance, the microbial diversity and metabolic interactions within these systems, particularly in understudied regions like the Brazilian margin, remain poorly resolved (5); (6); (7). Central to these ecosystems are methylotrophic microorganisms, which utilize reduced one-carbon compounds (e.g., methane, methanol) to fuel carbon, nitrogen, and sulfur transformations (8, 9). Methanotrophs, in particular, mitigate methane emissions and influence carbonate precipitation, yet their distribution and activity in non-seep deep-sea sediments are enigmatic (10). While amplicon sequencing studies in the Alpha Crucis Carbonate Ridge (ACCR) and adjacent Santos Basin have detected taxa such as *Methylomirabilota*, these approaches often fail to resolve low-abundance methanotrophs, leaving their functional roles speculative (5, 7). For instance, *Methylomirabilota* populations in ACCR sediments lack canonical methane oxidation genes, suggesting metabolic plasticity or methodological limitations in detecting rare taxa (6); (7). This knowledge gap underscores the need to explore the “rare biosphere”, i.e. microbial taxa present at low abundances but with potential outsized ecological impacts. Recent genomic advances reveal that rare microbes can harbor conserved adaptive traits, enabling persistence in oligotrophic environments(11);(12);(13). Such taxa may act as keystone species in carbonate-rich systems, maintaining metabolic functions during periods of resource scarcity. For example, reclassified genera like *Methylotuvimicrobium* (formerly *Methylomicrobium*) exhibit genomic adaptations to energy limitation, hinting at strategies for survival in different habitats (14). However, whether these taxa contribute to biogeochemical cycles without active seepage remains untested.

Here, we hypothesize that: (i) Aerobic methanotrophs constitute a cryptic component of the rare biosphere in deep-sea sediments under non-seep conditions, evading detection by standard amplicon sequencing; (ii) These organisms possess a conserved core genome optimized for oligotrophy, enabling persistence in low-energy environments; (iii) The rare biosphere’s metabolic versatility, particularly in deep-sea carbonate mound ecosystems, enables functional resilience, with aerobic methanotrophs sustaining carbon, nitrogen, and sulfur cycling through alternative metabolic pathways during periods of methane scarcity. To test this, we integrate culture-dependent enrichments, metataxonomics, and metagenomics of sediments cultures from the Santos Basin. By coupling these approaches, we aim to resolve the genomic and functional landscape of aerobic methanotrophs, assess their contributions to biogeochemical processes, and evaluate the ecological resilience of microbial consortia in deep-sea carbonate ecosystems.

## Materials and methods

### Study area and sample collection

Sediment sampling was conducted in the Santos Basin, located in the Southeast region of the Brazilian continental margin, occupying approximately 3.52 x 10^5^ km^2^. This basin covers the continental margin of the Southwest Atlantic and is limited to the north by Alto de Cabo Frio and to the south by the Florianópolis Platform. Samples were collected in November 2019 on board the R/V Alpha Crucis of the Oceanographic Institute of the University of São Paulo (IO-USP) during the development of the project Biology and Geochemistry of Oil and Gas seepages, SW Atlantic (BiOIL) (15). Sampling was developed in three distinct regions over the upper slope: (i) Area 1 - the Tupana Carbonate Ridge - TCR, a 35 km-long lineament of carbonate mounds with occurrence of Cold-Water Coral (CWC)(16), (ii) Area 2 - the Alpha Crucis Carbonate Ridge - ACCR, a ring-shaped carbonate ridge where there is evidence of seepage of hydrocarbons (4, 15) and (iii) Area 3 - an extensive pockmark field originated from salt tectonics, with the occurrence of an exhumed salt diapir - PF (1); (17) (Figure 1). Sediment samples were collected in triplicates using a stainless steel box-corer (50 cm x 50 cm) (Table S1). The sediment cores were sliced into 0-15 cm layers with sterile spatulas, and stored in evacuated Hungate tubes. The headspace was filled with argon and samples were preserved at 5 °C.

**Figure 1.**
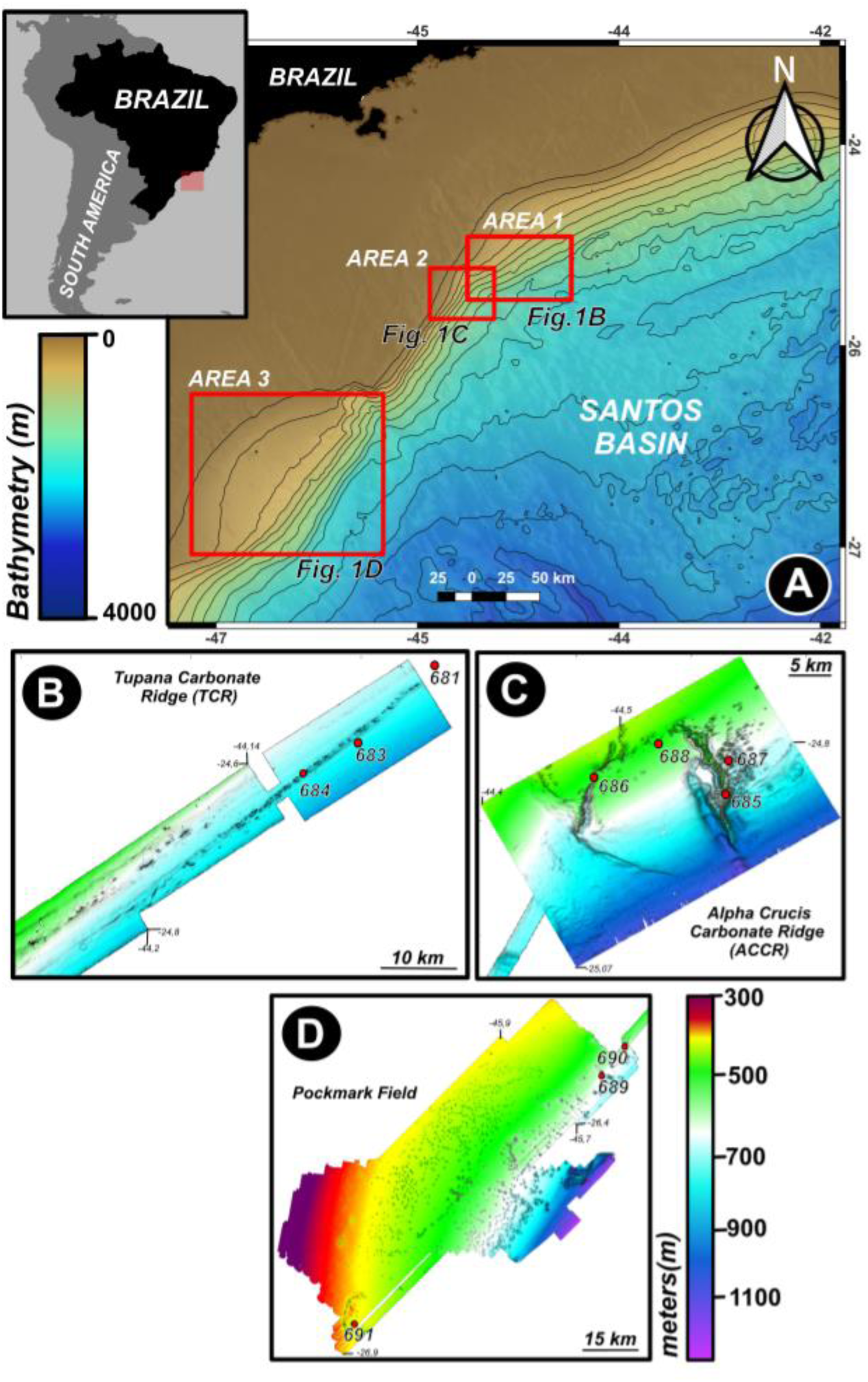
Bathymetric map of the southwestern Santos Basin, offshore southeastern Brazil, highlighting key geomorphological features. (A) Regional bathymetric overview indicating the locations of the three study areas: Area 1 (Tupana Carbonate Ridge – TCR), Area 2 (Alpha Crucis Carbonate Ridge – ACCR), and Area 3 (Pockmark Field). The inset map shows the location of the study area along the Brazilian margin. (B) Detailed bathymetric map from Tupana Carbonate Ridge (TCR) in Area 1. (C) Bathymetric view of the Alpha Crucis Carbonate Ridge (ACCR) in Area 2. Red dots represent sampling stations. (D) Bathymetric map of the Pockmark Field in Area 3. Color scales represent water depth (A) and seafloor elevation (B–D), with depth contours and bathymetric gradients indicating topographic variability. Coordinates are in decimal degrees.

### Cultivation of methanotrophic consortia

The 24 sediment samples were inoculated under aerobic and anaerobic conditions, yielding 48 enrichment cultures for methanotrophic cultivation. Aerobic and anaerobic cultures were performed in an NMS culture medium with copper addition (18) in BOD TE-371 (Tecnal, Brazil), at 20 °C. After autoclaving, the following were added: 10 mL, per liter of Phosphate Solution (26 g KH_2_PO_4_ and 62 g NaHPO_4_.7H_2_O in 1L of stock solution); 1.4 mg of CuCl_2_ 2H_2_O per liter of medium and 10 mL of sediment extract prepared from the deep-sea sediment obtained (Hamaki et. al., 2005). The headspace of the aerobic cultures (24 samples) was composed of methane:air (Whyte Martins, USA) in the proportion of 1:1, while the headspace in the anaerobic cultures (24 samples) was composed of methane:argon in the proportion of 1:1, totaling 48 cultures. During the suspension of laboratory activities in 2020 due to COVID-19, the cultures were kept in methane atmosphere, in an incubator BOD TE-371 (Tecnal, Brazil) at 5 °C to reduce microbial metabolism and favor the preservation of the consortia. The incubation period totaled 9 months. Methane concentrations in the headspace were monitored via gas chromatography (GC-FID) following Nakayama et al. (2014) (19), with calibrations using pure standards and filtered argon dilution to mitigate contamination.

### DNA extraction and sequencing of the 16S rRNA gene by Illumina Miseq

The DNeasy PowerBiofilm kit (MoBio, USA) was used for genomic DNA extraction, according to the manufacturer’s instructions. Approximately 15 mL of each culture was concentrated in a refrigerated centrifuge at 20 °C (12,000 x g for 15 minutes), and the precipitate was resuspended with the lysis solution. The extracted DNA was quantified using Qubit dsDNA HS Assay (Thermo-Fisher Scientific, USA). The 16S rRNA gene was sequenced using the Illumina Miseq platform, 2 x 350 bp, average coverage of 100,000 paired-end reads per sample, using universal primers 515F-Y 5’-GTGYCAGCMGCCGCGGTAA-3’, 926R 5’-CCGYCAATTYMTTTRAGTTT-3’ (Parada et al., 2015) at NGS Soluções Genômicas facility (Piracicaba, SP, Brazil). The sediment samples were extracted, sequenced and analyzed in a previous study (5).

### Bioinformatic analysis of the 16S rRNA gene

The 16S rRNA gene sequences were analyzed for their quality using the FastQC software. Bioinformatic analysis was performed using the QIIME2 software, v. 2020.2. Raw sequences were imported into QIIME 2 (v. 2020.2, https://docs.qiime2.org/) (5, 20) using the *qiime tools import* script via a manifest file. The sequences were summarized using the *qiime demux summarize* command and the DADA2 software was used to obtain the observed Amplicon Sequence Variants (ASVs) (21). Through the quality analysis generated by FastQC, both the forward and reverse sequences were truncated at position 240 and the barcodes were removed, using the *qiime dada2 denoise-paired script*. The taxonomy was signed using the *qiime feature-classifier* script using the SILVA v.138 database (21, 22). Phylogenetic distances were calculated using the script *qiime phylogeny align-to-tree-mafft-fasttree*, using the MAFFT aligner (23).

The alpha and beta diversity metrics were calculated using the *qiime diversity core-metrics-phylogenetic* script that calculates core diversity metrics. In the phyloseq package (v. 3.6.3) (23, 24) of R v. 3.6.3 (R Development Team, 2018) the Simpson and Shannon indices were calculated and visualized via box-plot. The non-parametric Kruskal-Wallis test was performed to determine whether there were significant differences in richness and evenness. Beta diversity was measured by the Bray-Curtis distance and visualized via NMDS (non-metric multidimensional scale). Permanova tests were performed to determine if there were differences in the dissimilarity of treatment applied to the cultures. To identify ASVs that are differentially abundant between treatments, the DESeq2 software was used, which performs differential expression analysis for sequence count data (25). The 16S rRNA gene sequencing data are available in the National Center for Biotechnology Information Sequence Read Archives under BioProject ID PRJNA1228598.

### Metagenomic sequencing of microbial consortia

Based on the results of metataxonomic analysis (16S rRNA gene), five consortia were selected for metagenomic sequencing, two consortia were from anaerobic cultures (_AN) and three were from aerobic cultures (_A): 685B1A, 686B3A, 684B1AN, 688B3AN and 683B3A. These consortia were filtered based on the results obtained in the 16S analysis. The metagenomic libraries were prepared using the Illumina DNAPrep Kit (Illumina, San Diego, CA, USA) and sequencing was performed in Illumina Hiseq platform (NextSeq2x 100p) at NGS Soluções Genômicas facility (Piracicaba, SP, Brazil).

### Recovering metagenome-assembled genomes (MAGs)

The samples were kept separate for the assembly and recovery of the MAGs, to avoid cross the combination of contigs from different consortia in the same MAG. Analyses were performed on the Kbase online platform (https://www.kbase.us/) (26). After the cutting step using the Trimmomatic v. 0.36 software (27) the metaSPAdes v. 3.15.3 software (28) was used for metagenomic assembly with a minimum contig length of 2,000 bp. The binning step was performed using three software: MaxBin2 v2.2.4 (29), CONCOCT v1.1 (30) and MetaBAT2 v. 2.3.0 (31). The consensus binning was performed using DAS-Tool v.1.0.7 (32), integrating results from MaxBin2, MetaBAT2, and CONCOCT. The quality of the resulting bins was assessed using CheckM v. 1.1.3(33). Bins were categorized according to the quality standards for MAGs proposed by Bowers et al. (2017) (34) as high-quality (completeness >90%, contamination <5%), medium-quality (completeness >50%, contamination <10%), and low-quality (completeness <50%, contamination <10%). All MAGs were taxonomically annotated using the GTDBTk v2.0.0 software (35). The annotation of contigs for the prediction of coding regions (CDSs) was performed using the RAST v1.073 (36) PATRIC (37) and PGAP(38), databases. The functional annotation of the MAGs was performed using the KEGG database using data from the DRAM (Distilled and Refined Annotation of Metabolism) tool v. 1.2.4 (39). Genes related to the methane, nitrogen and sulfur cycles were selected to be explored during the analyses.

### Comparative genomics between MAGs of the genus Methylotuvimicrobium

Two of the recovered MAGs were taxonomically classified within methanotrophic bacterial lineages. These genomes were selected for comparative genomic analysis with four publicly available methanotrophic genomes retrieved from the GenBank database (Table S2).

The tools ANI (Average Nucleotide Identity), AAI (Average Amino Acid Identity) and DDH (DNA-DNA in silico hybridization) were used to compare the MAGs and the reference genomes. The calculation of the ANI and AAI values was performed on the “Kostas Lab’’ website (http://enve-omics.ce.gatech.edu/g-matrix/) and the calculation of the DDH distance was performed on the genome-to-genome website (GGDC) (http://ggdc.dsmz.de/home.php). The pangenome analysis was performed with the anvi’o v.8 pipeline to compare reference genomes and MAGs, identifying the core region and singletons. The phylogenomic analysis was performed using the software Insert Genome Into SpeciesTree v2.2.0 (available in Kbase), which allows the construction of a species tree using a set of 49 universal core genes defined by families of COG genes (Clusters of Orthologous Groups) (40). Pangenome analysis was performed using the anvi’o v. 8 pipeline in order to conduct genome comparison and gene cluster identification. To determine the content of unique and shared genes among the genomes of methanotrophic microorganisms, clusters of orthologous genes were analyzed using the OrthoVenn3 program (41). The functional annotation of MAGs was performed using the KEGG database and the DRAM software (39) to search for complete metabolic pathways, comparison of genes of interest and search for methanotrophs genes.

## Results

### Comparing Aerobic and Anaerobic Cultures of Methanotrophs

Aerobic cultures presented 133 exclusive ASVs, while anaerobic cultures presented 112 exclusive ASVs and 133 were shared among all consortia (Figure S1). The microbial composition was similar between treatments in which *Proteobacteria* (92.6%) and *Bacteroidota* (7.29%) were the most abundant phyla (Figure S2). Other phyla such as *Firmicutes, Actinobacterota*, and *Bdelovibriota* accounted for less than 1% of the total abundance. The most abundant family was *Alcavoracaceae*, followed by *Pseudoalteromonadaceae, Sphingomonadaceae, Pseudomonadaceae* and *Marinobacteraceae*. The *Methylomonadaceae* family was also identified among the ten most abundant in enrichments (Figure 2A). Through the analysis of Beta diversity using the Bray-Curtis metric (Figure 2B), it was possible to identify that the samples are dissimilar regarding the treatment applied to the cultures. Dissimilarity was tested for statistical significance using a PERMANOVA (*Betadisper* p-value = 0.2788). The microbial composition of the aerobic cultures showed a significant difference (*p* = 0.001) compared to the anaerobic cultures, indicating that the applied treatment was responsible for generating a variation in the composition of the microbial communities.

**Figure 2.**
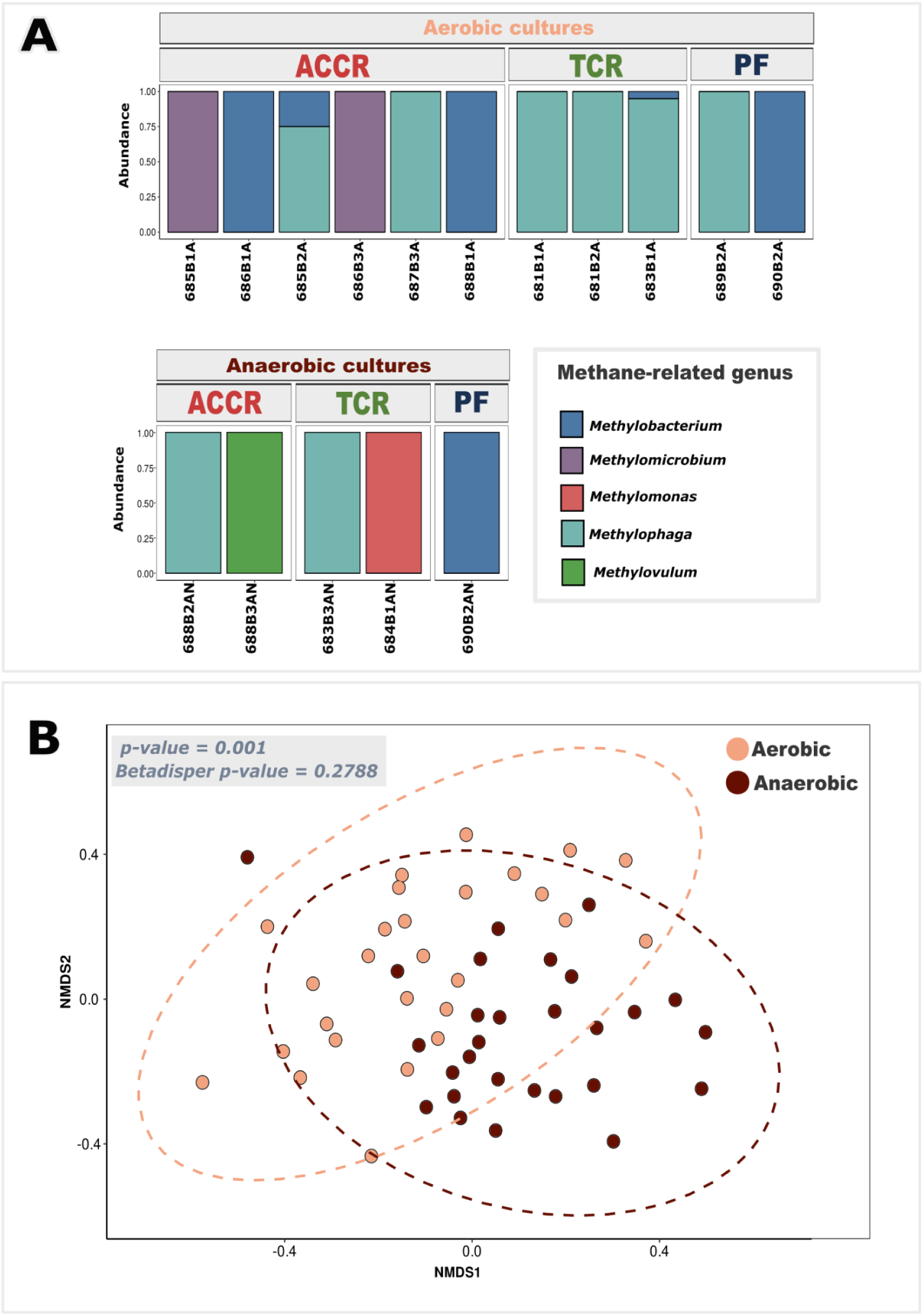
Composition and beta-diversity of methane-related genera in aerobic and anaerobic enrichment cultures from deep-sea carbonate systems. A. Relative abundance of methane-related genera (e.g., *Methylobacterium*, *Methylophaga*, *Methylovulum*, *Methylomicrobium*, and *Methylomonas*) in aerobic and anaerobic cultures from ACCR (Alpha Crucis Carbonate Ridge), TCR (Tupana Carbonate Ridge), and PF (Pockmark Field). B. Non-metric multidimensional scaling (NMDS) plot showing the distribution of aerobic and anaerobic samples based on the composition of methane-related microbial genera from ACCR, TCR, and PF. Statistical analysis: ANOSIM p-value = 0.001; Betadisper p-value = .2788.

The enrichment cultures from the Alpha Crucis Carbonate Ridge (ACCR), Tupana Carbonate Ridge (TCR), and Pockmark Field (PF) revealed three canonical aerobic methanotroph genera (*Methylomicrobium*, *Methylomonas*, *Methylovulum*) alongside facultative methylotrophs (*Methylobacterium*, *Methylophaga*) (Figure 2A). These taxa were undetected in original *in situ* sediment profiles (5), Figure S3), supporting their classification as rare biosphere members that proliferate under selective cultivation. Taxonomic composition diverged sharply with oxygen availability: *Methylobacterium* dominated aerobic cultures, whereas anaerobic incubations favored *Methylovulum* and *Methylophaga* (Figure 2B). NMDS ordination confirmed significant community dissimilarity (PERMANOVA, *p* = 0.001), with no dispersion bias (Betadisper, *p* = 0.2788), underscoring oxygen’s role in structuring methane-driven consortia.

Successful samples in the enrichment and recovery of MAGs from microorganism groups associated with methanotrophy refers to the collection sites 685 and 686, both located in the Alpha Crucis Carbonatic Ridge (ACCR) region. The Alpha Crucis carbonate mound was the region that presented the greatest diversity of genera of strict or facultative methanotrophic microorganisms (Figure 1B) when compared to Tupana Carbonate Mound and the Pockmark field. It was only possible to recover MAGs from methanotrophic microorganisms (*Methylotuvimicrobium*) from the ACCR probably due to the formation of this area of carbonate mounds.

### Metagenomic sequencing of consortia enriched with methanotrophic microorganisms and metagenome-assembled genomes (MAGs)

After sequencing the five metagenomes, 355.540.109 reads were generated (Table S3). The value of reads generated per sample can be seen in Table S3. The result obtained after assembly revealed that the number of contigs varied between 2053 and 6159, the percentage of GC content varied from 51.63% to 56.65% and the values of L50 and N50 varied from 102 to 408 and 151,495 to 46,552, respectively. All parameters analyzed in the assembly evaluation are recorded in Table S4.

We obtained a total of 90 MAGs, 78 high-quality MAGs, 8 medium-quality MAGs and 4 low-quality MAGs. The taxonomic annotation revealed the presence of 18 families of the Domain Bacteria, distributed in 3 different phyla. Seventy-four MAGs were taxonomically assigned as belonging to the phyla *Proteobacteria*, followed by 14 from the *Bacteroidota* and one from the *Actinobacteriota*, distributed in the classes *Gammaproteobacteria* (46 MAGs), *Alphaproteobacteria* (27 MAGs), *Bacteroidia* (13 MAGs) and *Actinobacteria* (1 MAG) (Figure S4). Additionally, two MAGs were taxonomically assigned to the genus *Methylotuvimicrobium*; both recovered from microbial consortia within the Alpha Crucis carbonate ridge called LECOM 001 and 002.

### Genomic analysis of MAGs from Methylotuvimicrobium sp. LECOM 001 and 002

Both MAGs showed a high completeness value, with 98.91% corresponding to LECOM 001 and 99.25% to LECOM 002 and low contamination values 2.53% and 2.5%, respectively. The recovered genomes are circular chromosomes of 5,020 Mb and (LECOM 001) and 5027 Mb (LECOM 002), with a content of 48.31% and 48.2% of GC content, respectively (Table 1). These values were similar when compared with the GC content of other genomes of the same genus, such as 48.6% of *Methylotuvimicrobium* sp. wino1, and the reference strains, *M. buryatense strain 5GB1C* (48.7%) and *M. alcaliphilum str. 20Z* (48.7%). Through the annotation, a total of 5,084 CDSs (Coding DNA sequences or DNA coding regions) corresponded to *Methylotuvimicrobium* sp. LECOM 001 and 5,109 CDSs to LECOM 002. The annotated hypothetical proteins totaled 2,183 and 2,206, respectively (Table 1).

**Table 1.**
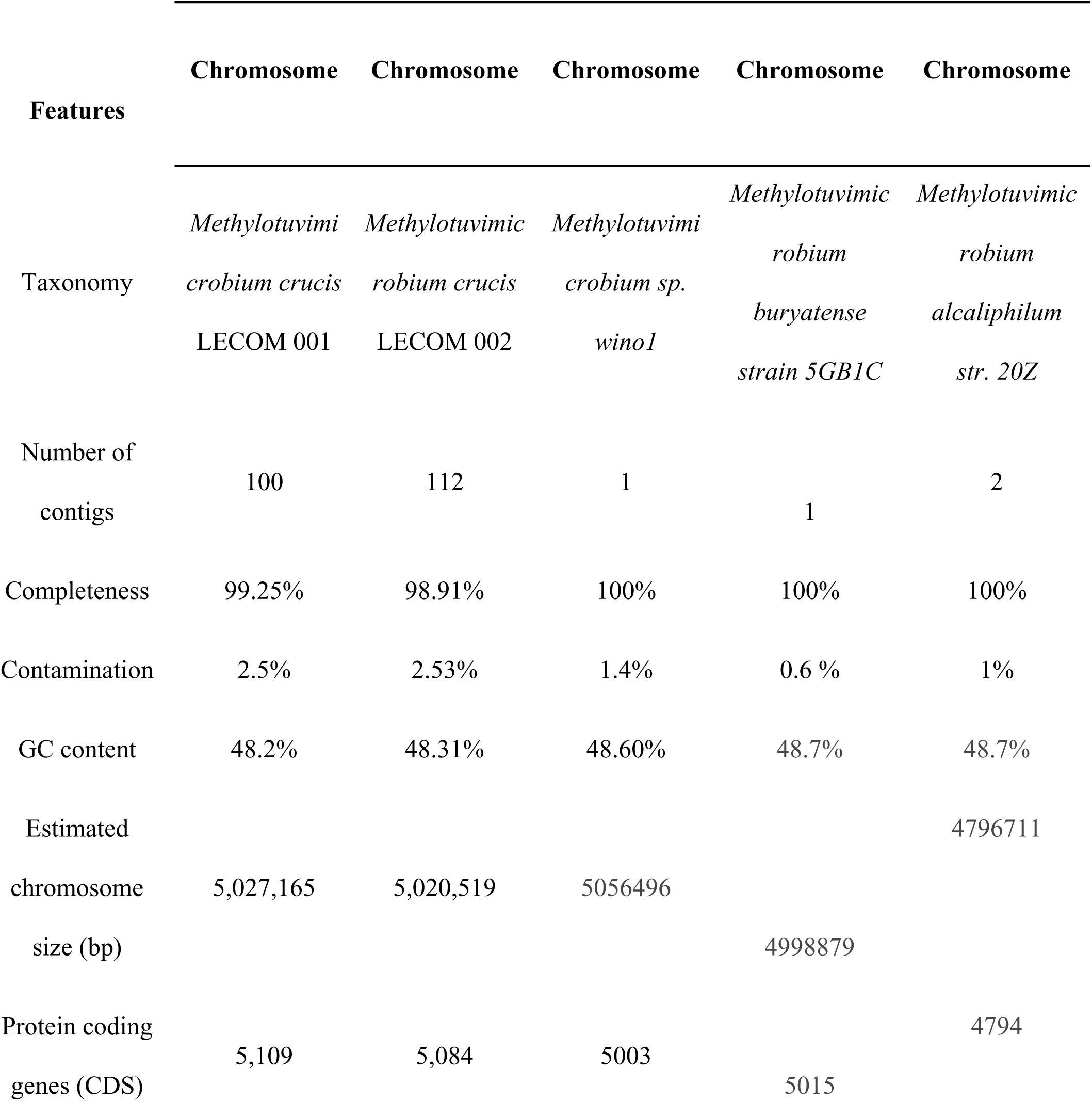
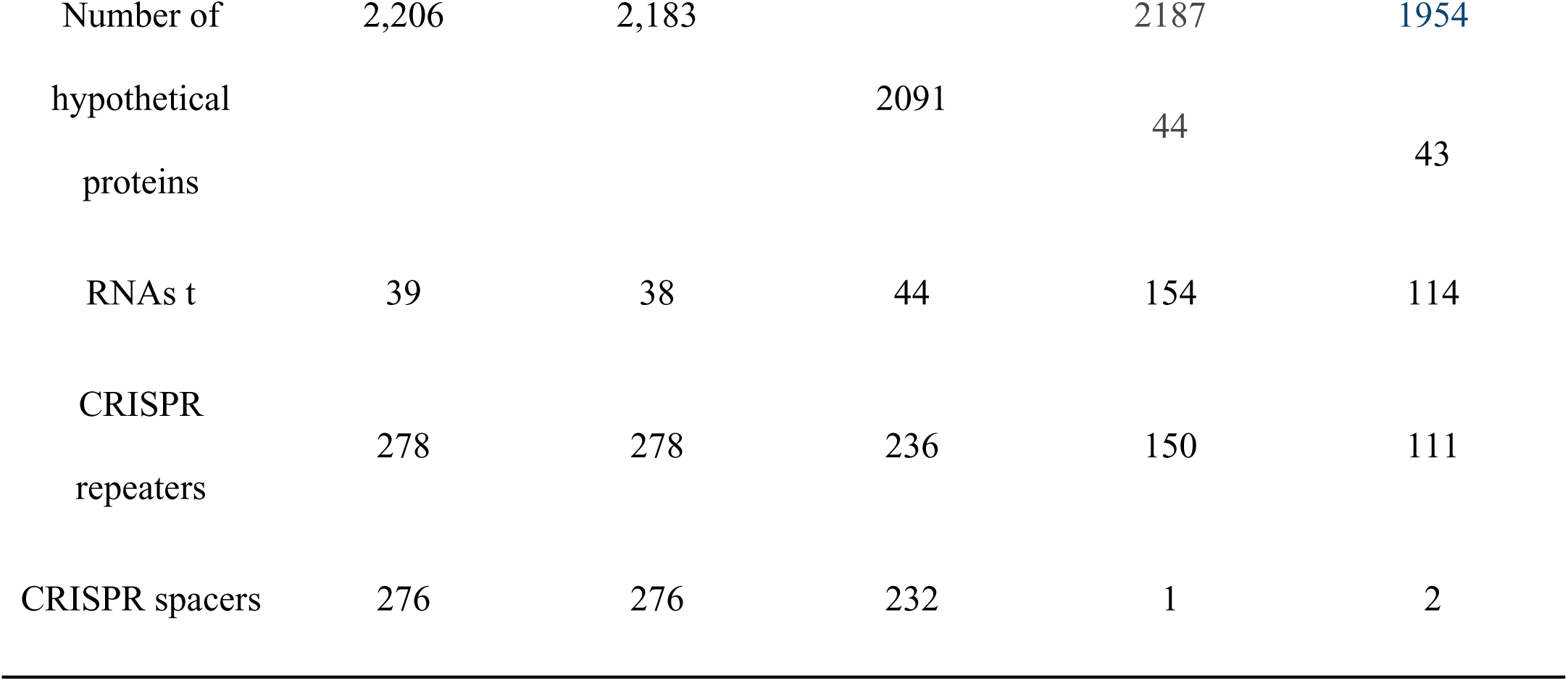
General features of the *Methylotuvimicrobium* LECOM 001, LECOM 002, wino1, *M. buryatense*, and *M. alcaliphilum* genomes. This table presents a comparative overview of key genomic features of *M. crucis* LECOM 001 and LECOM 002, alongside three reference genomes from GenBank: *M.* sp. wino1, *M. buryatense* 5GB1C, and *M. alcaliphilum* 20Z. Features include assembly statistics (number of contigs, estimated chromosome size), genome quality metrics (completeness, contamination), GC content, number of predicted protein-coding sequences (CDSs), number of hypothetical proteins, tRNA counts, and CRISPR system components (number of repeats and spacers).

The comparative analysis of overall genome-relatedness indices, including ANI, AAI and DDH showed that *Methylotuvimicrobium* sp. LECOM 001 and 002 correspond to the same species (Table 2). Following the current parameters for species delimitation, (Rodriguez & Konstantinidis, 2014) organisms considered to belong to the same microbial species share > 95% mean nucleotide identity (ANI), > 95% mean amino acid identity (AAI), and > 70% similarity (*in silico*) genome-to-genome hybridization (DDH).

**Table 2.**
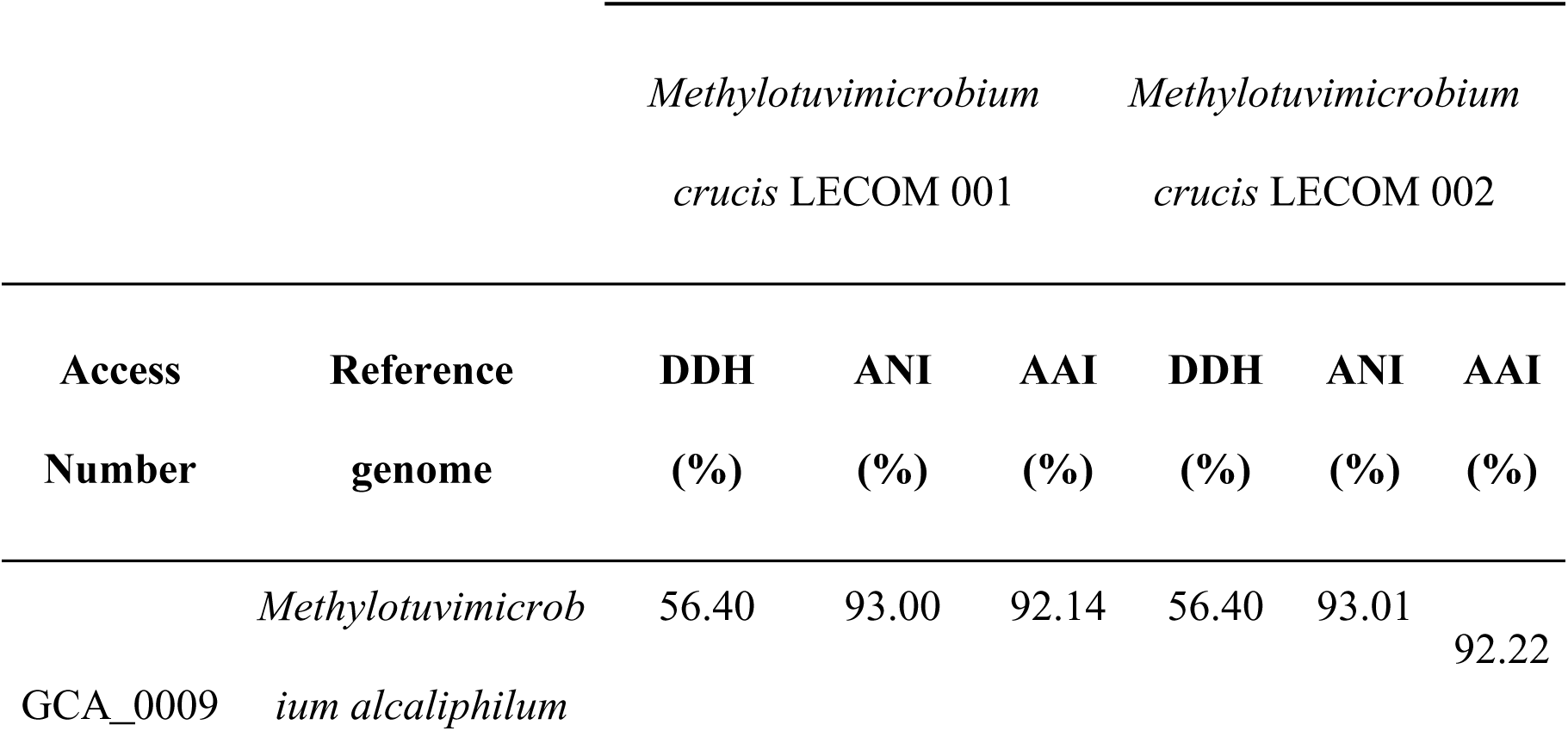
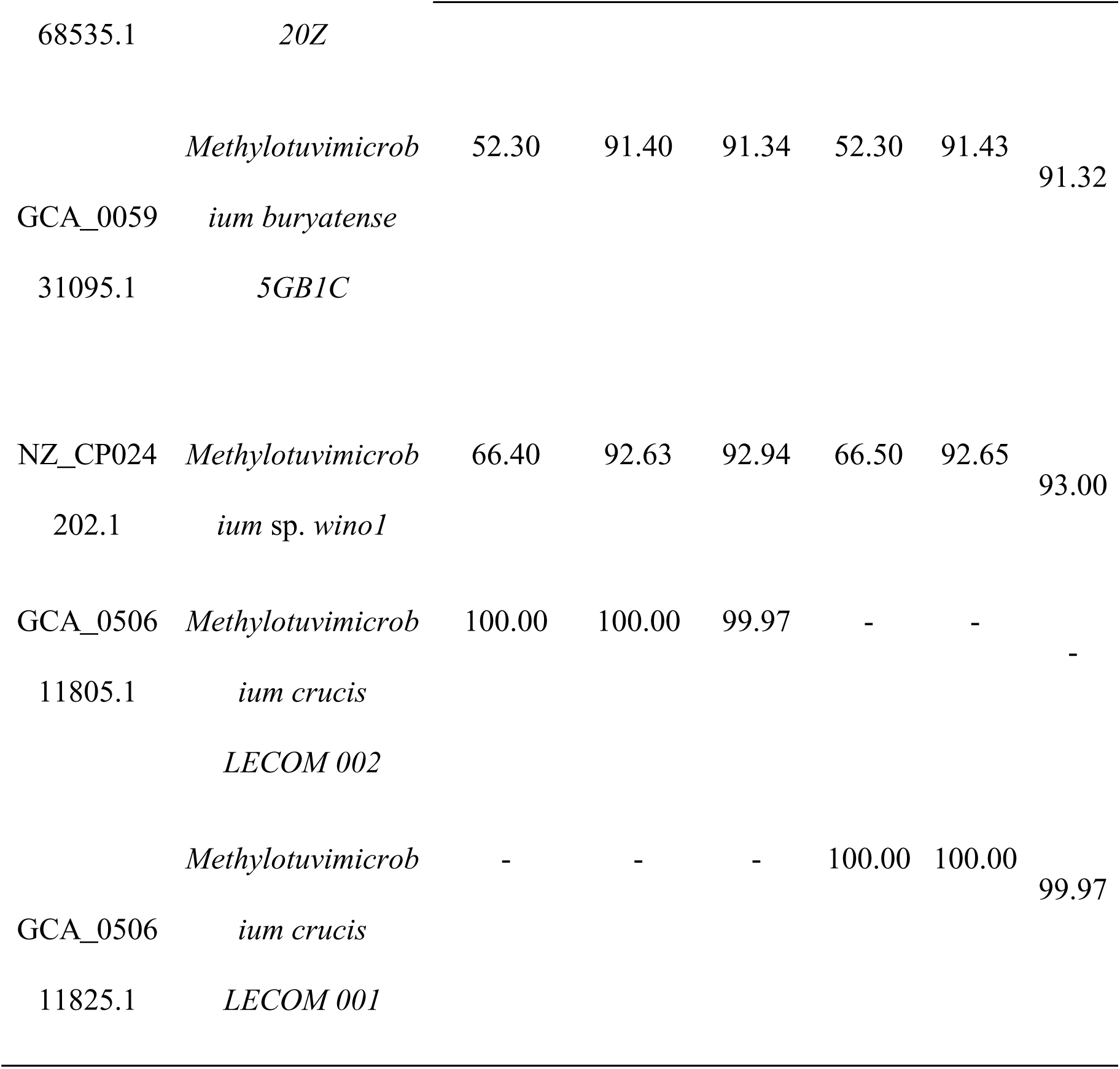
Genomic similarity metrics (DDH, ANI, and AAI) between *Methylotuvimicrobium crucis* strains LECOM 001 and 002 and reference genomes from GenBank. Values of DNA– DNA in silico hybridization (DDH), average nucleotide identity (ANI), and average amino acid identity (AAI) are shown for pairwise comparisons between M. crucis strains LECOM 001 and 002 and selected *Methylotuvimicrobium* reference genomes retrieved from GenBank.

The DDH, ANI, and AAI values indicate that the MAGs recovered in this study do not belong to any species of *Methylotuvimicrobium* available in Genbank. These results suggest that the MAGs found may represent new species of the genus, with genomic similarities to MAGs from marine environments but with unique characteristics.

Given these genomic distances, we propose that the MAGs LECOM 001 and 002 represent a novel species within the genus *Methylotuvimicrobium*, for which the name *Methylotuvimicrobium crucis* sp. nov. is proposed, according to SeqCode rules and recommendations (42). The specific epithet *crucis* refers to the Alpha Crucis carbonate mound, the site from which the genomes were obtained, and also to the Southern Cross constellation (*Crux*). Phylogenomic analyses support the monophyly of this taxon, which forms a well-supported clade with *Methylotuvimicrobium* sp. wino1, its closest known relative, while remaining genomically distinct based on all measured criteria (ANI, AII, DDH). Phylogenomic analysis revealed that the MAGs of *Methylotuvimicrobium* sp. LECOM 001 and 002 formed a clade with 100% bootstrap support, and three other MAGs from the same genus (Figure 3C). A distinct deep-sea clade was also observed, consisting of MAGs recovered from carbonate mounds in the Santos Basin and a MAG retrieved from deep-sea sediments. This phylogenetic grouping reinforces the similarities between the MAGs from deep-sea environments, suggesting a potential common adaptation to these ecosystems

**Figure 3.**
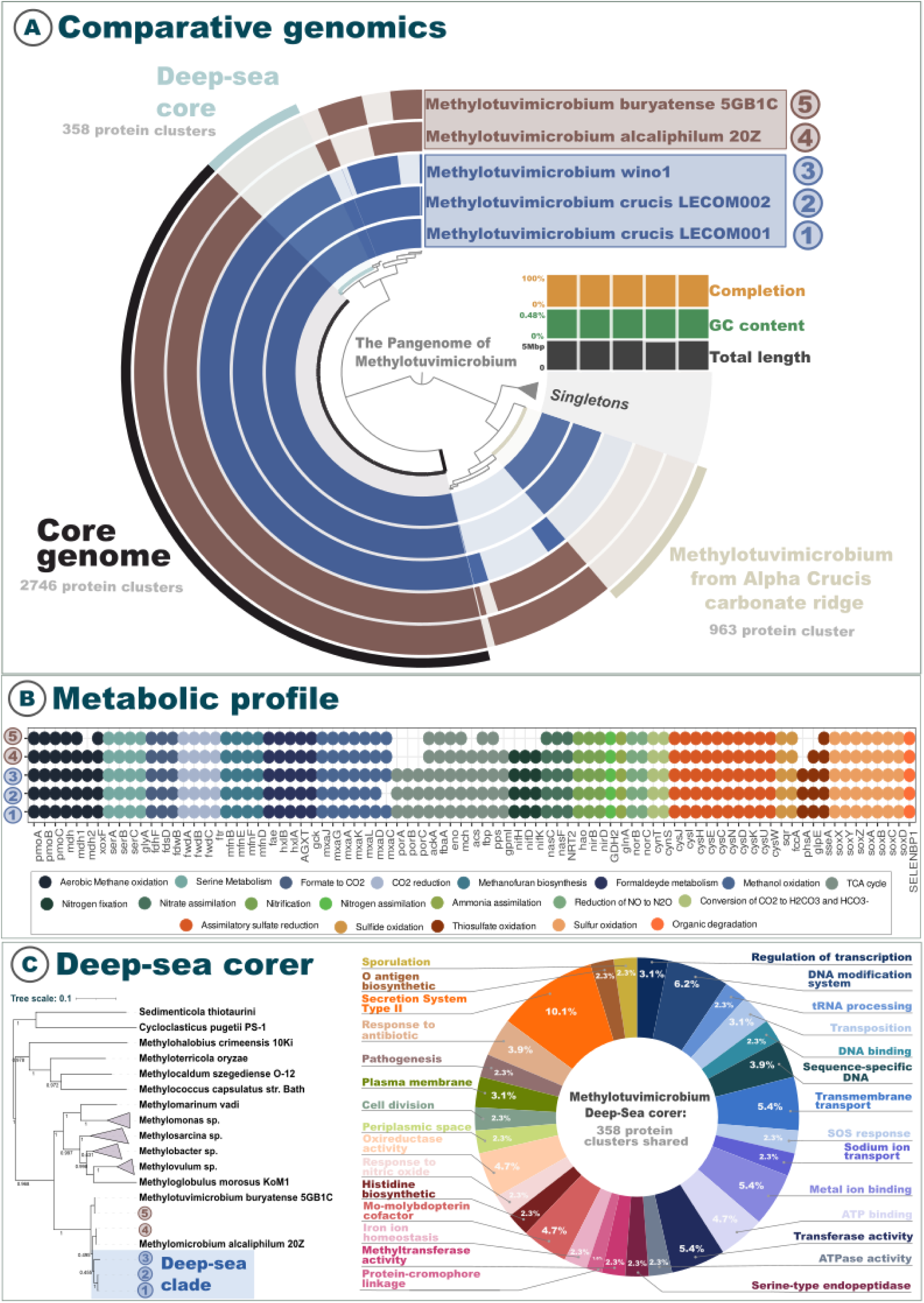
Comparative genomic and functional insights into deep-sea *Methylotuvimicrobium* lineages. (A) Comparative genomics: Pangenome analysis of *Methylotuvimicrobium* strains, including two newly recovered genomes from deep carbonate mounds (LECOM001 and LECOM002), compared with reference genomes. The core genome, shared by all strains, comprises 2,746 protein clusters, while a subset of 358 clusters defines the *deep-sea core*, exclusive to the deep-sea lineage. Additionally, 983 protein clusters are unique to a strain recovered from the Alpha Crucis carbonate ridge. The inner ring includes metadata for each genome, including completeness, GC content, and total genome length. Singleton genes (unique to a single genome) are also indicated. (B) Metabolic profile: Presence/absence matrix of key metabolic pathways across *Methylotuvimicrobium* genomes, inferred from functional gene annotations. Pathways are grouped by metabolic categories, including methane oxidation, nitrogen and sulfur transformations, carbon fixation, amino acid biosynthesis, central carbon metabolism (e.g., TCA cycle), and stress response mechanisms. Colored dots represent the presence of genes or pathways, highlighting functional differentiation among deep-sea and non-deep-sea strains. (C) Deep-sea core gene functions: Left panel shows a maximum-likelihood phylogenetic tree, highlighting the deep-sea clade (blue). The right panel presents the functional classification of the 358 protein clusters that compose the deep-sea core genome. Functional categories include transcriptional regulation, DNA modification, stress response, cell adhesion, metal ion binding, and horizontal gene transfer-related functions, underscoring potential adaptations to deep-sea environmental pressures.

Pangenome analysis comparing recovered MAGs with reference MAGs identified a core region among the five genomes belonging to the same bacterial genus, a region that includes 15342 genes (Figure 3A). The pangenome analysis of the genus *Methylotuvimicrobium* revealed a core set of genes essential for the survival and adaptation of these species (Table S6). Key genes in the core genome include those associated with aerobic methane oxidation, carbon fixation, cellular structure, intracellular transport and signaling, and DNA processing genes such as endonucleases. Several general functions were identified, including intracellular trafficking, secretion, vesicle biogenesis, translation and ribosomal structure, cell cycle control, cell wall and membrane biogenesis, and the transport and metabolism of nucleotides, carbohydrates, lipids, amino acids, and cofactors, highlighting the genus’s metabolic complexity. Stress-related genes include cold-shock proteins from the *CspA* family and *CheY-like* domains, which are involved in cold responses, chemotaxis, and sporulation regulation. Bacterial motility is supported by genes involved in the biosynthesis and functionality of flagella and type IV pili, including components such as *FliH, MotA, MotB, PilZ, PilU*, and *FlgC*, which are essential for movement and environmental interaction. The pangenome also features a robust DNA repair system, with genes involved in SOS response (*LexA*), UV damage repair (*PhrB*), and proteins such as *UvrD* and *MutS* that participate in mismatch repair and recombination. Genetic mobility is evidenced by transposases, prophage elements, and plasmid maintenance systems such as *VapI*, emphasizing the genomic plasticity of the genus. Additionally, genes related to chemotaxis and signal transduction, such as *CheW, CheA*, and *CheB*, were identified, enabling these bacteria to sense and respond to environmental chemical gradients.

### The marine core

In addition to the core region, we identified a deep-sea core (1234 genes) containing regions shared only among deep-sea *Methylotuvimicrobium*, including the two recovered MAGs and the *Methylotuvimicrobium*sp. wino1 genome (Figure 3A).

Our findings suggest that *Methylotuvimicrobium* from deep-sea environments exhibit strong adaptive mechanisms through horizontal gene transfer and biofilm formation. For instance, the genomes harbor genes associated with the mobilome, including prophages and transposons, contributing to genomic adaptability. Key transposases identified include the *TnpA* family, *YbfD/YdcC*-associated transposase, and inactivated *TnpA* derivatives. The plasmid stabilization protein *ParE* was annotated, suggesting mechanisms for plasmid stability in challenging environments. Phage-related proteins, such as *gpD* (phage protein D), *FI* (phage tail sheath protein), and *gpI* (P2-related tail formation protein), indicate the presence of prophages or remnants, potentially influencing host fitness. Furthermore, proteins involved in pilus formation, such as *PilF/PilW* (Type IV pilus assembly proteins) and *TadD* (Flp pilus assembly protein), emphasize the role of pili in bacterial adhesion and biofilm formation. These findings suggest that *Methylotuvimicrobium* from deep-sea environments exhibit strong adaptive mechanisms through horizontal gene transfer and biofilm formation. We also observed proteins involved in DNA repair and maintenance, including *RpnC* (recombination-promoting DNA endonuclease) and *Nfi* (deoxyinosine 3’-endonuclease), essential for repairing oxidative DNA damage and maintaining genomic stability under the high-pressure, low-temperature conditions of the deep-sea. Additional proteins, such as *Ssb* (single-stranded DNA-binding protein) and *Mod* (adenine-specific DNA methylase), suggest mechanisms for DNA stabilization and protection against environmental stressors.

Transcriptional regulation is crucial for managing oxidative stress and metabolic flexibility. Regulators such as *lysR, csgD*, and *merR* (*SoxR*) control gene expression in response to environmental signals, while sigma factors like *rpoE* (σ24) and helix-turn-helix (HTH) domain regulators (*yiaG* and *copY*) fine-tune metabolic pathways. Essential cofactors, such as molybdenum (via *moaA*, *moaD, moaE*) and cobalamin (via *btuB*), are synthesized, supporting survival in nutrient-limited environments. Enzymes like *hisF, hisH*, and *gadA* contribute to nitrogen metabolism. For cellular trafficking and secretion, *Methylotuvimicrobium* utilizes the Type II secretory pathway, with components like *gspA, exeA*, and *gspD/PulD*. *Flp* pilus assembly proteins (*tadD, tadG*) and the *vgrG* protein are involved in surface attachment and biofilm formation.

### Functional annotation of MAGs

Functional annotation of genes allowed observing which genes are related to important biogeochemical cycles in the environments in which these microorganisms are present. Comparing the five MAGs of the genus *Methylotuvimicrobium*, we were able to observe that there are metabolic differences when comparing the deep-sea group (LECOM 001, LECOM 002 and wino1) with the group from other environments. The pangenome and the analysis of orthologous proteins made it possible to identify a core genome that is shared only between *M*. wino 1, *M*. LECOM 001 and *M*. LECOM 002 (Figure 3A), revealing a highly versatile metabolic capacity in relation to C₁ compound utilization, central metabolism, and nitrogen/sulfur cycling.

For aerobic methanotrophy and ammonia oxidation, genes for particulate methane monooxygenase activity *pmoA-amoA, pmoB-amoB*, and *pmoC-amoC* are consistently present across all strains (*Methylotuvimicrobium crucis* LECOM 001, *Methylotuvimicrobium crucis* LECOM 002, *Methylotuvimicrobium* sp. wino1, *Methylotuvimicrobium buryatense* strain 5GB1C, *Methylotuvimicrobium alcaliphilum* str. 20Z), indicating the widespread capacity of these strains to oxidize methane aerobically.

For methylotrophy, nearly the full *mxa* gene cluster (*mxaFJG ACKLD*, except *mxaI*) is present across all strains, which encodes for calcium-dependent methanol dehydrogenase (*MDH*), showing a robust ability to metabolize methanol into formaldehyde. Notably *mxaC* is missing in *Methylotuvimicrobium* crucis LECOM 002, potentially indicating a functional divergence in methanol dehydrogenase assembly or activity. Other related genes, such as *mdh1* and *xoxF* are also present in all strains, exhibiting different forms of *MDH*. The gene *mxaI* is only present in *Methylotuvimicrobium alcaliphilum* str. 20Z, suggesting slight variations in methanol oxidation pathways.

Downstream processing of formaldehyde is supported by the presence of genes *fae*, *hxlB*, *hxlA*, *AGXT*, and *gck*, which mediate formaldehyde detoxification and assimilation via the ribulose monophosphate and serine pathways (*serABC*, and *glyA*). The same consistency is observed in genes involved in formate to CO_2_ conversion (*fdoG*, *fdhF*, *fdwA*, *fdsD*, *fdwB*) and CO_2_ reduction (*fwdA*, *fmdA*, *fwdB*, *fmdB*, *fwdC*, *fmdC*, *ftr*) via the Wood–Ljungdahl pathway, highlighting the organism’s capacity to thrive in fluctuating redox conditions. Genes associated with methanofuran biosynthesis (*mfnB, mfnE, mfnF, mfnD)* are present in all strains, indicating a strong capacity for methanofuran production, an essential component in methanogenesis.

Core carbon metabolism is supported by complete TCA cycle components such as *porA, porB, porC,* and *porG*, along with genes involved in gluconeogenesis and the Embden-Meyerhof-Parnas pathway, including *ackA, fbaA, eno, gpmI, mch, acs, fbp, pps*, and *ppsA*. This suggests robust energy generation and carbon flux capacity across diverse substrates. Notably, *porA, porB, and porC, porG* are absent in *Methylotuvimicrobium buryatense* strain 5GB1C and *Methylotuvimicrobium alcaliphilum* str. 20Z, indicating potential differences in TCA cycle functionality or alternative metabolic pathways.

The genes *nifHDK* for nitrogen fixation are present in *Methylotuvimicrobium crucis* LECOM 001, *Methylotuvimicrobium crucis* LECOM 002, *Methylotuvimicrobium* sp. wino1, and *Methylotuvimicrobium alcaliphilum* 20Z, indicating their capacity for nitrogen fixation, a key process for converting atmospheric nitrogen into a biologically usable form. However, these genes are absent in *Methylotuvimicrobium buryatense* 5GB1C, suggesting that this strain may not be capable of nitrogen fixation under natural conditions and might rely on other nitrogen sources.

Genes for nitrite and nitrate assimilation and dissimilation were identified, including *nasC, nasA, nrtA, nasF, cynA, NRT2, narK,* and *nrtP* are present in all genomes, suggesting they can utilize nitrate and cyanate as nitrogen sources in limited ammonium availability. In addition, the presence of *hao, nirB*, and *nirD* across all strains supports the potential for nitrite reduction and hydroxylamine oxidation associated with dissimilatory nitrogen metabolism linked to energy conservation under low-oxygen conditions. Additionally, denitrification capability was detected by the presence of *norB* and *norC.* Finally, genes *cynT* and *cynS* involved in reaction of cyanate with bicarbonate to produce ammonia and CO_2_ are present in all strains. These features confirm a conserved mechanism across all strains to support a flexible nitrogen metabolism suited for fluctuating redox and nitrogen-limited habitats for both nitrogen uptake and reduction.

Sulfur metabolism is similarly well represented across all strains. For assimilatory sulfate reduction, several key genes, including *cysU, cysW, cysD, cysH, cysIJ*, and *cysE,* are present in all genomes. These genes play crucial roles in the reduction of sulfate to sulfide. Regarding oxidation of reduced sulfur compounds, genes such as *sqr* and *fccA* are also present in all genomes. Sulfide oxidation is a critical process for converting hydrogen sulfide (H₂S) into elemental sulfur or sulfate, contributing significantly to sulfur cycling. Sulfur oxidation involves several genes, such as sox system genes (*soxX, soxY, soxZ, soxA, soxB*, *soxC, and soxD*) (Zhou et al., 2025). Among these, *soxX, soxA*, and *soxB* are conserved across all strains. Additionally, the gene *SELENBP1*, associated with organic sulfur compound degradation, is present in all strains, suggesting that these bacteria can degrade organic sulfur compounds, and an additional layer of redox adaptability.

## Discussion

### Rare but Resilient: The Ecological Role of Aerobic Methanotrophs in Deep-Sea Carbonate Systems

Our study provides evidence that aerobic methanotrophic bacteria, although underrepresented in standard amplicon sequencing, are important components of the rare biosphere in carbonate mounds and pockmark areas. Only after using enrichment techniques was it possible to detect the presence of some methanotrophic groups, including *Methylomicrobium*, *Methylomonas*, and *Methylovulum*, which were not detected in the initial analyses. This difference highlights the limitations presented by culture-independent methods in detecting low-abundance taxonomic groups and the importance of combining culture-dependent and culture-independent techniques to reveal specific microbial groups.

Furthermore, the results suggest that aerobic methanotrophic groups remain components of the rare biosphere, potentially contributing to nutrient cycling in the sediment even in the absence of methane seeps, but can respond rapidly to episodes of methane escape. Our study reinforces the relevance of studying regions such as carbonate mounds and pockmark areas to better understand the aerobic methanotrophic groups that make up the rare biosphere and their ecological role in these regions.

### Genomic Adaptations and Metabolic Versatility of Deep-Sea Methanotrophs

The conserved genomic content shared between the deep-sea *Methylotuvimicrobium* strains (LECOM 001, LECOM 002, and wino1) distinguishes these lineages from their shallow-water relatives, reflecting adaptations to high pressure, low temperature, and oligotrophic conditions. This genomic core underscores the metabolic versatility that enables these organisms to thrive in extreme environments. This metabolic versatility is supported by various genomic features that reflect specific adaptations to deep-sea conditions (43). The environment imposes severe physiological constraints, including high hydrostatic pressure, low nutrient availability, and oxidative stress (44); (45); (46). To endure these challenges, the *Methylotuvimicrobium* MAGs encode multiple stress-resistance mechanisms, including DNA repair systems *(RpnC, Nfi, UvrD*) to counteract pressure-induced damage, cold-shock proteins (*CspA*) to stabilize protein function at low temperatures, and heavy metal resistance genes (*CopZ, CzcD*) to manage trace metal scarcity. The presence of sulfur assimilation pathways (e.g., *cysJI*, *sqr*) suggests an ability to respond to fluctuating redox conditions in sediment environments. Together, these features mirror those found in other piezophilic microbes and point to a conserved deep-sea survival toolkit that enables persistence in this extreme yet ecologically significant habitat.

Another striking feature of the deep-sea *Methylotuvimicrobium* MAGs is the enrichment of mobile genetic elements (e.g., *TnpA*, *ParE*) and prophage remnants (*gpD*, *FI*), suggesting that horizontal gene transfer (HGT) contributes significantly to their adaptation ability. The presence of Type IV pilus assembly proteins (*PilF*, *TadD*) and cyclic di-GMP signaling domains (*GGDEF*, *EAL*) further points to biofilm formation as a core survival strategy, likely facilitating attachment to sediment particles and enhancing access to limited nutrients (47); (48). These traits are consistent with those observed in other deep-sea microorganisms, where genetic exchange and surface adhesion are crucial for persistence in energy-limited environments (3). The genomic features of these *Methylotuvimicrobium* strains, spanning stress tolerance, metabolic flexibility, and ecological interactions, may reflect a finely tuned adaptation to the deep-sea niche. Their capacity for methane oxidation, nitrogen fixation, and sulfur metabolism positions them as potential keystone taxa in deep-sea biogeochemical cycles. Rather than transient or passive microbial community members, these organisms appear to be specialized residents shaped by genomic streamlining and functional redundancy. While their ecological significance is becoming clearer, important questions remain about the *in situ* regulation of these pathways and the extent to which they influence deep-sea productivity. Targeted approaches combining high-pressure cultivation with metatranscriptomics will be key to unlocking these dynamics and refining our understanding of their roles in global nutrient cycling. The genomic multifunctionality of *Methylotuvimicrobium crucis*, spanning sulfur assimilation (*sox, sqr*), denitrification (*norBC*), and nitrogen fixation (*nifHDK*), challenges the paradigm of methanotrophs as methane-dependent specialists. Instead, their metabolic plasticity positions them as keystone taxa capable of sustaining elemental cycling in energy-limited carbonate mounds, even without active seepage. The Alpha Crucis Carbonate Ridge’s geological setting, marked by inferred methane migration (4), suggests these methanotrophs act as latent methane filters, potentially mitigating emissions during episodic fluxes. Their nitrogen-fixing capacity mirrors diazotrophic methanotrophs in cold seeps (49), hinting at a role in priming nutrient-poor sediments for microbial colonization. The observed genomic adaptations in the *M. crucis* lineages such as biofilm formation (GGDEF/EAL domains), mobile elements, and stress-response genes potentially support resilience to oligotrophy and fluctuating methane availability. Collectively, the genomic features highlight the deep-sea core potential for methane oxidation, formaldehyde assimilation, full respiratory and fermentative carbon metabolism, flexible nitrogen and sulfur cycling, and biofilm formation as ecological adaptations for persistence and competitiveness in redox-variable systems among substrate pulses, which is critical for survival in dynamic deep-sea habitats.

Future studies should prioritize *in situ* activity assays, such as stable isotope probing of methane assimilation or nitrogen fixation rates, to resolve their contributions to carbon sequestration and microbial food webs. Additionally, quantifying their influence on carbonate precipitation could clarify linkages between methanotrophy and long-term carbon burial in non-seep systems.

## Conclusions

This study uncovers the hidden ecological role of active deep-sea methanotrophs in the rare biosphere of SW Atlantic carbonate-rich sediments. By integrating cultivation with genomics, we propose *Methylotuvimicrobium crucis* sp. nov., a novel obligate methanotrophic species represented by metagenome-assembled genomes (MAGs) from the Alpha Crucis carbonate ridge, the first methanotroph genomes reported for marine sediments in this underexplored region. Strikingly, these methyl-dependent taxa thrive in an area without active methane seeps, challenging paradigms about their survival in energy-limited habitats. Genomic evidence further reveals their potential for nitrogen fixation, suggesting a dual role in methane and methanol oxidations and nutrient cycling that could sustain microbial consortia in deep-sea carbonate ecosystems.

This discovery highlights a critical blind spot in marine microbial surveys: functionally specialized taxa like *M. crucis* evade detection in standard metabarcoding studies due to their low abundance yet may drive biogeochemical processes in cryptic niches. The presence of a putative new *Methylotuvimicrobium* species in SW Atlantic carbonates raises fundamental questions: Are transient methane pulses or alternative energy sources subsidizing these communities? Does their metabolic plasticity enable persistence during methane scarcity?

Future studies should target in situ activity assays to validate the proposed nitrogen fixation capability and methane metabolism under dynamic conditions. Additionally, systematic exploration of SW Atlantic cold seeps and carbonate-hosted methane reservoirs is urgently needed to resolve the mismatch between methanotroph occurrence and detectable methane fluxes. By bridging cultivation with omics, this work expands the genomic diversity of *Methylotuvimicrobium*. It also advocates for a unified framework to decode the ecological strategies of microbial “dark matter” in deep-sea sediments.

As the first genomic record of methanotrophs in SW Atlantic carbonates, *M. crucis* exemplifies the untapped functional diversity lurking in geologically complex marine regions, a frontier for understanding microbial resilience and its role in mediating Earth’s methane balance.

### Data availability

Raw amplicon-sequencing reads and MAGs recovered from metagenomic data are available on NCBI’s Read Archive under accession PRJNA1234549. The register of *Methylotuvimicrobium crucis* sp.nov. can be found at the following link https://seqco.de/i:49843.

## Author Contributions

ACAB: investigation, methodology, formal analysis, visualization, writing - original draft, Writing - review & editing.

FMN: investigation, methodology, writing - original draft, Writing - review & editing.

FVP: investigation, methodology,writing - original draft, Writing - review & editing.

FMS: investigation, methodology, writing - original draft, Writing - review & editing.

AGB: investigation, methodology, writing - review & editing.

RB: investigation, methodology, visualization, writing - original draft, Writing - review & editing.

MMM: funding acquisition, project administration, writing - review & editing.

PYGS:funding acquisition, project administration, writing - review & editing.

VHP: funding acquisition, supervision, writing - original draft, Writing - review & editing.

## Funding

This research was carried out in association with the R&D project registered as ANP 21012-0, “MARINE LIFE - BMC - OIL, AND GAS SEEPS (BIOIL)” (Universidade de São Paulo / Shell Brasil / ANP) – “Biology and Geochemistry of Oil and Gas Seepages, Southwest Atlantic (BIOIL)”, sponsored by Shell Brasil under the ANP R&D levy as “Compromisso de Investimentos com Pesquisa e Desenvolvimento”. The BIOIL Project (Shell Brasil) supported the following fellowships: BIOIL Project (Shell Brasil) supported the ACAB Master’s fellowship and AGB and FMN postdoctoral fellowships.

## Conflict of Interest Statement

The authors declare that the research was conducted without any commercial or financial relationships that could be construed as a potential conflict of interest.

## Acknowledgments

We thank the research teams of LECOM and Rosa C. Gamba for their invaluable support. Rosa was a dedicated technician in our laboratory, whose commitment and generosity left a lasting impact on our team and this work. This manuscript is also a tribute to her memory. We also thank the captain and the crew of the R/V Alpha Crucis (IO-USP, FAPESP Process number 2010/06147-5) for the essential support during the I BIOIL and II BIOIL oceanographic cruises.

